# A machine learning based analysis to probe the relationship between odorant structure and olfactory behaviour in *C. elegans*

**DOI:** 10.1101/2021.07.26.453815

**Authors:** Aayushi Vishnoi, Rati Sharma

## Abstract

The chemical basis of smell remains an unsolved problem, with ongoing studies mapping perceptual descriptor data from human participants to the chemical structures using computational methods. These approaches are, however, limited by linguistic capabilities and inter-individual differences in participants. We use olfactory behaviour data from the nematode *C. elegans*, which has isogenic populations in a laboratory setting, and employ machine learning approaches for a binary classification task predicting whether or not the worm will be attracted to a given monomolecular odorant. Among others, we use architectures based on Natural Language Processing methods on the SMILES representation of chemicals for molecular descriptor generation and show that machine learning algorithms trained on the descriptors give robust prediction results. We further show, by data augmentation, that increasing the number of samples increases the accuracy of the models. From this detailed analysis, we are able to achieve accuracies comparable to that in human studies and infer that there exists a non trivial relationship between the features of chemical structures and the nematode’s behaviour.

## I. INTRODUCTION

Olfaction is one of the primary senses that helps an organism navigate its environment, i.e., search for food, avoid potential threats and even find mates [1]. The study of olfaction, therefore, is one of the major research areas and model organisms such as *Caenorhabditis elegans* have been used since the early 1990s to elucidate the relationship between chemical environments and related animal behavior [2]. *C. elegans* is a free-living nematode that has been a preferred model organism not only for olfaction studies, but also for various other research areas, ranging from aging to behaviour, ever since it was first used by Sydney Brenner in the 1960s [3]. This is primarily due to the presence of genes and signaling pathways that have homologs in humans and the relative ease in performing genetic manipulations.

The *C. elegans* olfactory system, itself being a constituent of the larger chemosensory system, helps it integrate and make use of ecologically relevant information efficiently [4–6]. Its chemosensory system consists of 32 neurons that are either directly or indirectly exposed to the environment and helps sense both soluble and volatile (olfactory) cues, with AWA, AWB and AWC being the most prominent ones among them. The soluble chemicals or odorants bind to the receptors present on these neurons, and the downstream signalling these trigger give rise to various behavioral phenotypes.

The first assays to observe the behaviour of worms in response to chemicals were carried out by Ward in 1973 [7], followed by Dusenbery in 1974 [8]. However, later in the 1990s, there were much more elaborate studies carried out by Bargmann et al. [9, 10] which not only described worm behaviour to a long list of chemicals but also discovered the associated neurons and genes involved. These studies also gave way to an interest in getting insights into the dynamics of worm movements during chemotaxis [11, 12], with a large number of works now focusing on finding the accurate mathematical model that can describe the same [13]. Since then, there is a growing list of known *C. elegans* attractants and repellents, which has lead to efforts in harnessing the olfactory behaviour to aid diagnostics in tuberculosis [14] and cancer [15]. The individual or group of neurons involved in the sensing have been identified using ablation [9, 16] and RNAi studies [17]. Additionally, sensory glia cells have also been implicated in olfaction [18, 19], albeit in a neuron independent manner.

The olfactory representation of molecules in the brain, however, remains an unsolved problem even with a considerable amount of work being put into deciphering the underlying structures and mechanisms involved. With research on vision being aided by machine learning approaches, there is now an interest in applying computational techniques to olfaction research as well, both in human and animal studies. There have been efforts to construct an odorant chemical space and connect it to the resulting subjective experience of smell [20] through deep learning [21, 22] and supervised machine learning [23] approaches using data from human subjects. Alternatively, there is also an interest in learning the organisation of olfactory systems to make better deep and machine learning algorithms as opposed to the pre-existing ones that derive inspiration from the organisation of visual processing systems. Most of the work in this area has been in insects, with a relatively simpler olfactory system when compared to humans, and this has resulted in the development of olfaction based classification [24] and similarity search algorithms [25].

Most of the studies concerning human olfaction use the Dialogue for Reverse Engineering And Methods (DREAM) olfaction challenge dataset [20], which suffers from the drawback of the perceptual descriptors being subjective and highly variable between individuals. This is because of genetic variability, which manifests in the form of variability in olfactory receptors and, in turn gives rise to differences in olfactory perception [26]. Apart from genetics, odor perception depends on age, environmental factors and linguistic capabilities of individuals [27]. However, that is not the case with lower animals, especially in the isogenic population of *C. elegans* grown and maintained in a laboratory setting, which are genetically identical and are also exposed to similar environments.

With a simple nervous system and its connectome being complete[28], *C. elegans* can be useful in building biologically inspired neural networks, as was done recently by Hasani et al. [29]. It’s simple nervous system nonetheless gives rise to complex behaviours, which renders it an ideal system to study the problem of olfactory representation and construct simpler models for the same. In that light, a recent study employs computational approaches to predict the structures of olfactory receptors in *C. elegans*[30]. The authors used protein threading followed by a simulated molecular docking approach to find out the relationship between the structure of odorant receptors present on the neurons and the chemicals that bind to it. Therefore, in this work, we curate experimental data from the literature and propose a machine learning approach to find out if there are underlying relationships between *C. elegans* olfactory behavioural data and odorant structure. Our proposed method for the olfaction-chemical structure problem is based on observables rather than subjective experience. It is employed here to perform a binary classification task linking olfactory behaviour to chemical structure and can easily be tested for reliability using standardised behavioral assays.

The rest of the article is organized as follows. Section II gives an overview of the approach followed and the models analyzed. Section III presents the results obtained and compares the accuracies of the various models. The final section, Section IV, provides a discussion of the results and reflects on possible future extensions of this study.

## II. METHODS

The complete research scheme is shown in Fig. 1. The key steps are as follows: (i) Data curation and cleaning, (ii) Pre-processing to extract features in the form of molecular descriptors or one-hot encodings, (iii) Model training and (iv) Model validation. The following sections go into details of each of these four steps.

**Figure 1:**
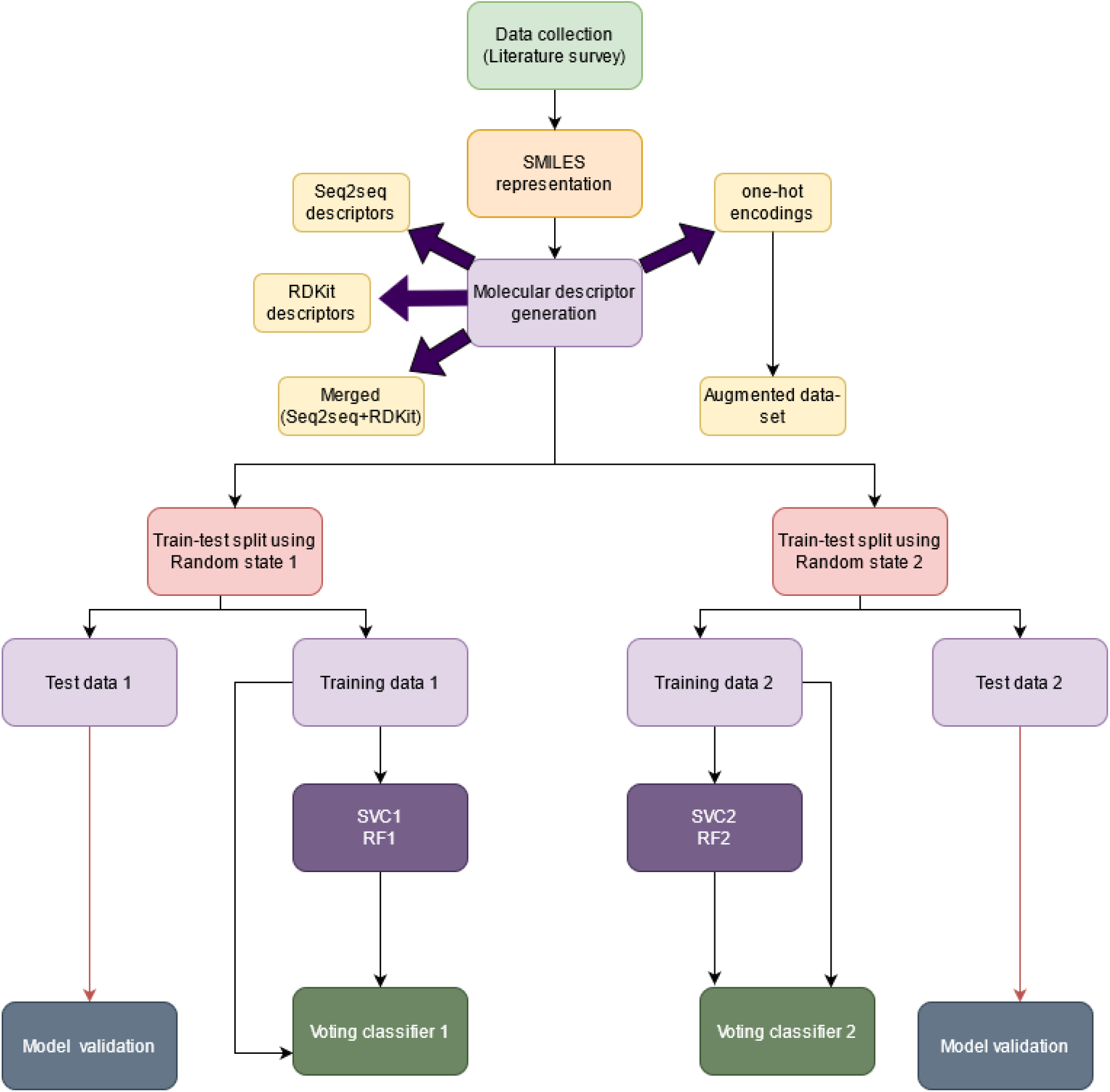
Flow chart showing the steps involved

### A. Data set preparation

#### 1. Data collection

The first step in analyzing the structure-behavior relationship is data curation. To this end, we carried out a literature survey to make a list of chemicals with known corresponding *C. elegans* behaviours [5, 6, 9, 10, 12, 14, 31–55]. However, different concentrations of some of the chemicals have been reported to show conflicting *C. elegans* behaviours. Therefore, we removed such chemicals from the list in order to avoid any inadvertent discrepancies. The resulting data set consists of 192 chemicals, 100 of which are attractants, 44 are repellents, and the rest are neutral. Since the number of attractants far exceeds the number of repellent and neutral chemicals, we assigned the chemicals to either one of the two classes - “attractive” or class “1” (100 chemicals) and “not attractive” or class “0” (92 chemicals) - in order to get a more balanced data set.

The labels (or classes) assigned to the chemicals are a result of the observations made through various experimental chemotaxis assays perfomed on *C. elegans*. However, due to the heterogeneity in the methods used among various studies, especially in terms of differences in concentrations and tests, the precise values of the quantitative measures are not considered, and instead, qualitative information is used for analysis. The most commonly used assay in the literature, for example, is the chemotactic index assay, wherein, a single or multiple worms are allowed to move freely on a plate that has both the control and the test chemical. The Chemotactic Index (CI) is then given by the fraction of worms that prefer the test chemical over the control. CI is therefore calculated using the following equation.

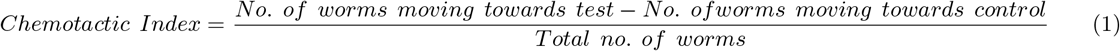

The chemicals showing significant positive CI values (> 0.1) are assigned to the class “attractive”, and the rest, which either have negative values or values that are not significantly different from 0, are assigned to the class “Not attractive”. This list is provided in the Supplementary Information, Table S1.

#### 2. Pre-processing

After having collected and labeled the data, the next step in the process is feature extraction. In Quantitative structure-activity relationship (QSAR) studies, such as those used in drug discovery and toxicology research, molecular descriptors are used as features for the purpose of applying algorithms to the data [56, 57]. These descriptors are generally symbolic or mathematical representations of chemicals that allow a computational analysis to be carried out on them. They can either be obtained from experimental measures of physical and chemical properties or can be calculated theoretically from different multi-dimensional representations of chemicals-for example, fragments [58], SMILES (2D) and graphs (3D) [59]. Depending on the method used, they can either have a direct physical interpretation or can be abstract representations of the properties encoded without direct interpretations. Simplified molecular input line entry system (SMILES) is one such system that represents molecules in the form of strings of characters. This system can be thought of as a language with simple vocabulary and defined grammatical rules. It is also a convenient way to store chemical information and has been used for the prediction of drug-target interactions using machine learning methods [60]. SMILES representations of molecules have also been used recently in a human olfactory prediction study [22] that employed deep neural networks. In our study, these representations were generated using the Chemical Identifier Resolver service designed by the CADD group of the Chemical Biology Laboratory at the National Cancer Institute [61].

The next step in pre-processing is to extract molecular descriptors from the SMILES representation of the chemicals. One such method developed by Winter et al. uses deep learning methods to learn a latent vector based representation of molecules that can accurately translate them from one format (SMILES) to the other (InCHI) [62]. This tool, in turn, is built on Xu et al.’s work on Seq2seq methods in natural language processing [63].

Similar methods have also been used by Gomez-Bombarelli et al. that allow the exploration of the chemical space of molecules to find their neighbours with applications in drug designing [64]. For the present study though, Winter et al.’s software was found more suitable as it allows extraction of molecular descriptors at an intermediate step [62]. However, having been designed for drug discovery, there is a pre-processing step involved in the tool that removes out salts and inorganic molecules. The resulting data set, therefore, only has organic molecules (with stereo-chemical information removed) with the following properties: molecular weight between 12 and 600 Da, more than 3 heavy atoms, and partition coefficient (measure of the solubility of a compound in two immiscible liquids at equilibrium), log P, between 7 and 5. Further analysis was carried out with the remaining 173 organic compounds - 81 belonging to class 0 and 92 to class 1. Molecular descriptors were also generated by RDkit using the ChemDes platform [65]. The resulting data set had 189 entries-99 belonging to class 1 and 90 belonging to class 0. Further, the two types of descriptors, viz. (i) from Winter et al.’s Seq2seq tool and (ii) from RDkit, were merged, keeping the molecules that were common to both data sets. This data set had 170 entries-90 belonging to class 1 and 80 to class 0. The models were trained on each of the three data sets and checked for prediction accuracy.

### B. Machine learning models

#### 1. Model selection

After having obtained the features in terms of molecular descriptors for the data set, machine learning (ML) methods were applied to it in a python setting. In earlier studies, especially those involving problems in a biological setting, two ML methods have been used extensively - Support Vector Classifier (SVC) and Random forest (RF) [66–68]. SVC operates by constructing a hyperplane to distinguish between classes, with the form of the hyperplane being determined by the underlying kernel and other parameters. Random forest, on the other hand, is an ensemble model that picks up decision trees that are fit to the data at random and uses the majority voting to give its predictions. Both of these are robust algorithms that learn from the data and can be used for better fits via hyperparameter optimizations. In our study, we used both these models and their variations in order to find the best model for the task of binary classification of the data set.

Before training the models though, the data set itself was first split into training and test sets in a 70-30 ratio using the train_test_split() function of the sci-kit learn library. For the train-test split, a random integer seed is used to shuffle and split the data. It is not a hyperparameter to be tuned, however, at times models are known to be overfitted to a particular random state. In order to avoid that, in our workflow, the same procedures were repeated for two different random states. To further avoid biases that might result during splitting, the stratify parameter was set to TRUE. This made sure that the same proportion of the two classes was maintained in both training and test sets.

Following the above mentioned train-test split, Python’s sci-kit learn library was used to fit simple random forest and SVC models, the results of which are shown in later figures. Apart from SVC and RF, other machine learning (ML) algorithms such as Gradient Boosting Classifier(GBC) and K-Nearest Neighbors (KNN) were also applied to the data in a similar manner. Since the initial data set consisted of a large number of features (more than twice the samples), Principal component analysis (PCA) was applied to it in order to avoid the curse of higher dimensionality. PCA is a dimensionality reduction technique that takes the projection of data from a higher dimensional space onto a lower dimensional space such that the highest variance comes to lie along the first principal component, the second highest along the second principal component and so on, with all the principal components being orthogonal to each other. The threshold for percentage of variance explained was kept at 80 percent and optimum number of principal components were then selected for each case via trial and error. The results of the PCA fitted models are given in Table S2.

PCA assumes the relationship to be linear which might not always be the case so we also tried non linear dimensionality reduction techniques, but they performed worse than the PCA models. The python implementation of the automatic feature selection method, Boruta (BorutaPy) was also tried for Random Forest models. It works for ensemble methods and creates shadow features according to a given threshold. It then compares them with real features and selects the ones that perform better at the task than the shadow features. This helps in reducing the number of features while keeping the original features rather than some linear or non linear combination of them as is the case in other dimensionality reduction methods.

We finally also used a *Voting Classifier* to make the best prediction. A voting classifier trains on various models and predicts the output by considering the majority vote from them. It can employ two types of voting strategies - hard voting and soft voting. In hard voting, the predicted output is the class which has the highest number of votes from the different models, while in soft voting, an average of prediction probabilities from the various models is taken, and the class with the highest average probability is selected as the output predicted class. To further increase the accuracy and to do away with any individual shortcomings the standalone models might have, two ensemble models were made using the voting classifier for both the random states. The confusion matrices and ROC-AUC curves for the same are given in Figs.2(d) and 3(b), respectively, more context for which will be provided in the next Section.

**Figure 2:**
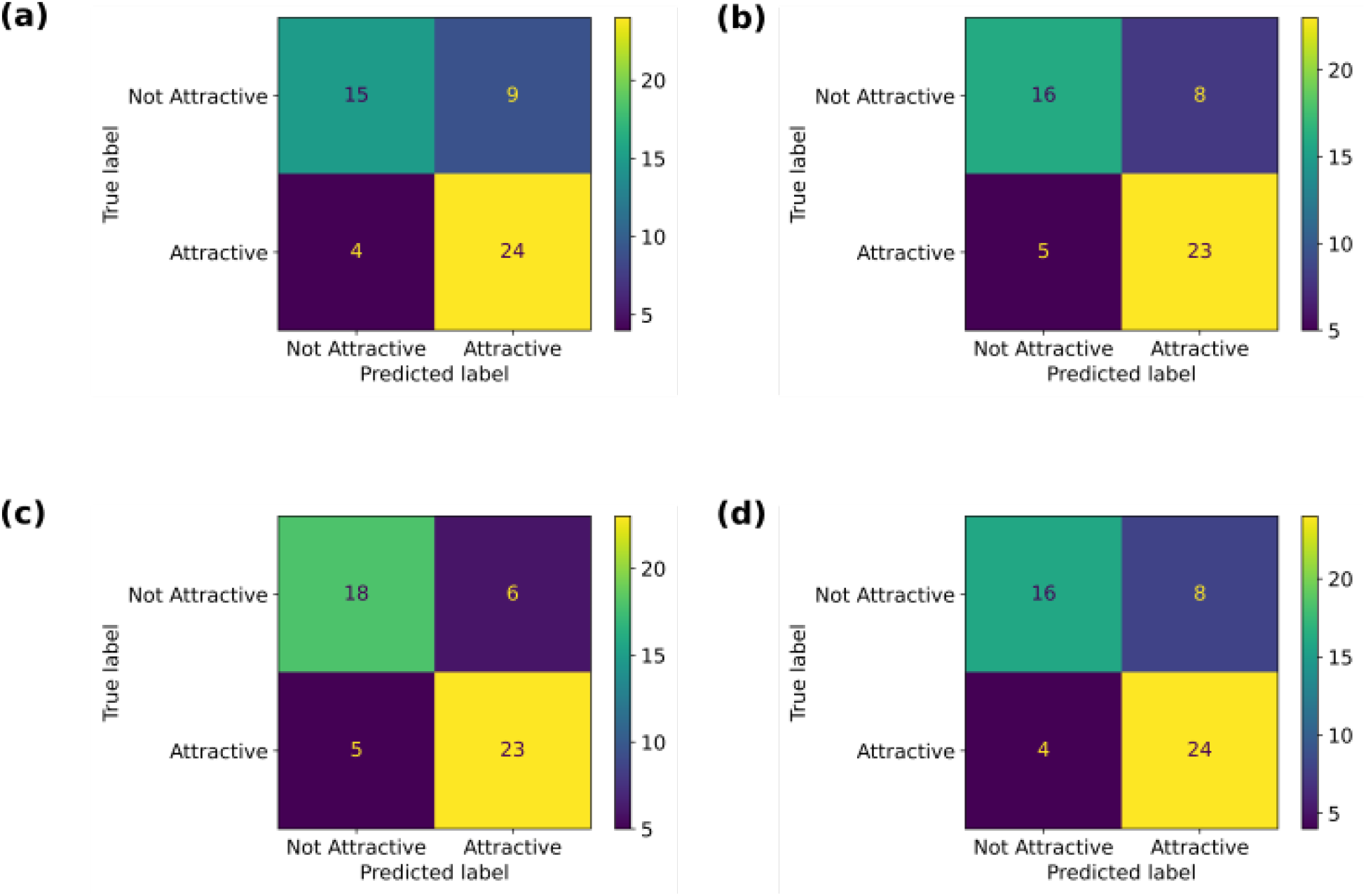
Confusion matrices for the first random state for. (a) Support Vector Classifier (SVC), (b) Random Forest (RF), (c) Gradient Boosting Classifier (GBC) and (d) Voting Classifier

**Figure 3:**
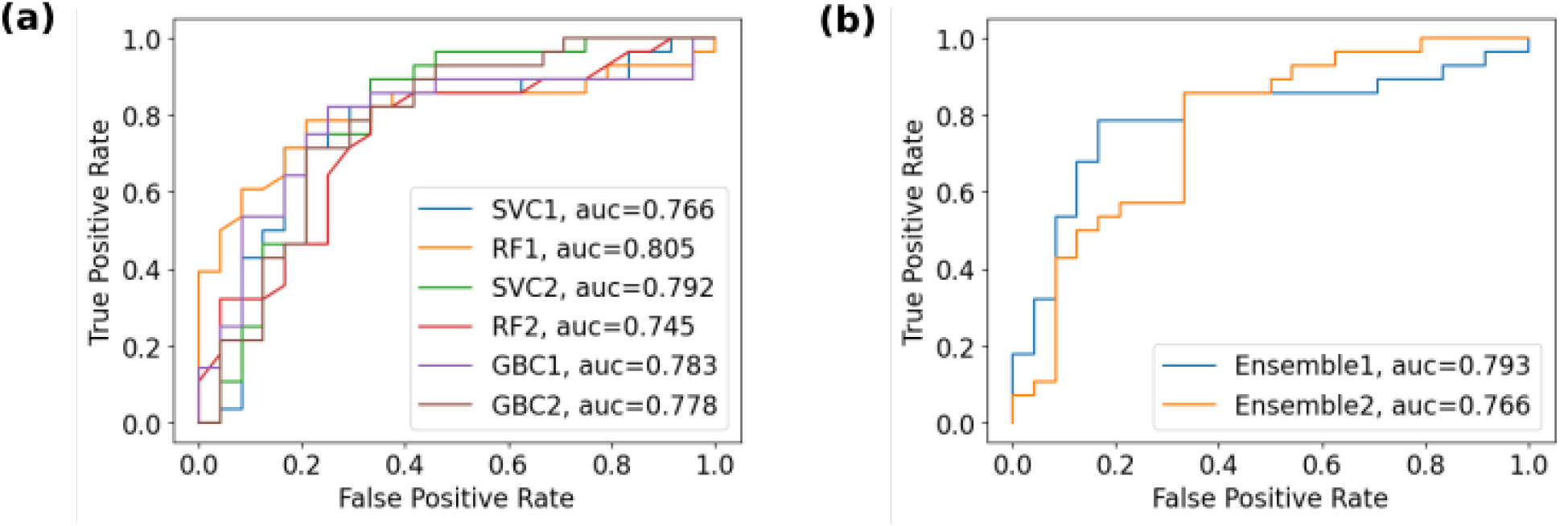
AUC-ROC curves for the following models trained using Seq2seq descriptors: (a) Support Vector Classifier in the first random state (blue) and the second random state (green), Random Forest in the first random state (orange) and the second random state (red), Gradient Boosting Classifier in the first random state (purple) and the second random state (brown); (b) Ensemble model in the first random state (blue) and the second random state (orange)

#### 2. Hyperparameter optimization and performance evaluation

We carried out a five-fold cross-validation and used a grid search to optimize the hyperparameters of the SVC models. However, similar hyperparameter optimization process was not carried out for Random forest models since the algorithm has inbuilt mechanisms for the same.

The following performance metrics were used for the purpose of evaluating the models:

1. **AUC-ROC scores:** The area under the receiver operating characteristic curve (AUC-ROC) is a performance metric used in binary classification problems to assess the ability of the classifier to distinguish between the two classes. It gives the probability of separating into the classes at various threshold values by plotting True Positive Rate (TPR, also known as recall) against False Positive Rate (FPR), where TPR and FPR are given by:

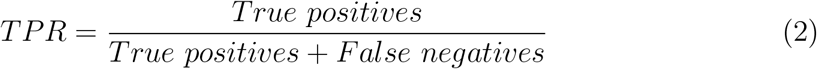

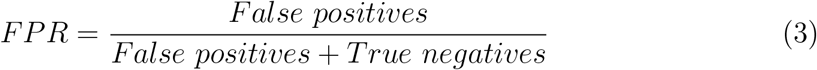 The curves were obtained for SVC and RF models for both the random states. The model with a steeper curve (higher positive slope) represents higher probability of a true positive in comparison to a false positive and therefore gives a better AUC score. The curves along with the AUC scores for these four models are given in Fig.3.
2. **F1 scores:** F1 score is a performance metric which can be described as a weighted sum of Precision and Recall and is given by

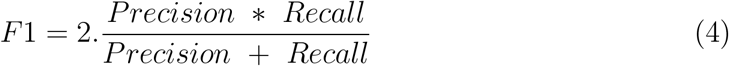 With,

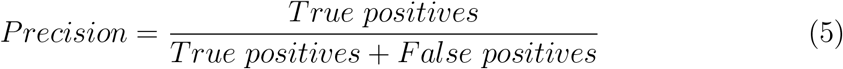

and Recall defined in Eq. 2

#### 3. Data augmentation

Data augmentation can solve the limited data availability problem and has been carried out for image data sets in the past [69, 70]. Since then, similar efforts have also been made to augment datasets for QSAR studies [71–73] and have been shown to increase the accuracy of prediction. One of these augmentation methods developed by Bjerrum [71] makes use of the fact that chemicals can be seen as graphs with the atoms being nodes and the bonds being the vertices of the graph. The whole graph can then be traversed starting from any node in the graph. Doing so gives rise to different SMILES strings for the same molecule and in effect increases the data set available for training. This method utilizes the randomization of SMILES using the RDKit library of python, followed by vectorization, which gives rise to one-hot encodings on which machine learning algorithms can be trained.

The average number of atoms present in the molecules for the data set (containing 173 molecules) was calculated to be 10, and that was used as a factor to augment SMILES. This was done in order to prevent biasing the data set in the favour of molecules that have a higher number of atoms. The resulting data set contained 11X (original smiles + 10X) the number of entries. After the duplicates were removed, the final data set consisted of 1404 entries-692 belonging to class 0 and 712 belonging to class 1. Further, Random Forest, SVC and other models were trained on the original as well as the augmented data following the same process as described for Seq2seq descriptors.

## III. RESULTS

### A. Behaviour is dependent on the chemical signature of the odorant

As stated in SectionII of this article, we converted the list of chemicals into three different types of feature sets, namely, (i) Seq2seq, (ii) RDKit and (iii) a combination of Seq2seq and RDKit descriptors, which we call Merged descriptors. We trained and tested each of these three descriptors on SVC, RF, KNN and a combination of these. Fig.2 shows the confusion matrices for the various models in the first random state trained using the Seq2seq descriptors. The respective performances of the standalone models are comparable to that of the ensemble model (Voting Classifier) of SVC and RF, with GBC performing the best (11 mis-classifications) out of these.

The AUC-ROC curves, along with the corresponding AUC scores for the standalone and ensemble models trained using Seq2seq descriptors in two random states are given in Figs.3(a) and 3(b) respectively. These too give comparable scores for all the models with one of the models, GBC applied to random state 1, giving the best score of 0.807.

A selection of the models tested are listed in Table I, while the complete set is listed in Table S2. Table I also summarises the F1 scores for various models obtained after dimensionality reduction and hyperparameter tuning, in both random states and for the three different descriptor sets. Barring one discrepancy in the case of the SVC model for RDKit descriptors, all the other models gave comparable predictions in both the random states. Following are the main observations from this analysis.

- For Seq2seq descriptors, the models SVC, RF, KNN and GBC give F1 scores that are comparable to each other as well as to the ensemble models that considered voting from RF and SVC. GBC performed the best among the standalone models followed closely by SVC and RF.
- In the case of RDKit descriptors, ensemble models of SVC and RF perform better than the individual models, but there is an overall reduction in scores when compared to that trained on Seq2seq descriptors. Here too, GBC performs best among all the models.
- Overall reduction in scores is also seen in the case of merged descriptors with GBC models performing better than the rest.

**Table I:** F1 scores of various models

| Model | Random state 1 | Random state 2 |
| --- | --- | --- |
| **Seq2seq Descriptors** |  |  |
| Support vector classifier | 0.787 | 0.750 |
| Random Forest | 0.780 | 0.762 |
| Random Forest(BorutaPy) | 0.733 | 0.7 |
| KNN | 0.764 | 0.767 |
| Ensemble(RF+SVC) | **0.800** | 0.800 |
| Gradient Boosting Classifier | **0.807** | 0.794 |
| **RDKit Descriptors** |  |  |
| Support vector classifier | 0.767 | 0.638 |
| Random Forest | 0.716 | 0.689 |
| Random Forest(BorutaPy) | 0.706 | 0.708 |
| KNN | 0.698 | 0.656 |
| Ensemble (RF+SVC) | 0.758 | 0.698 |
| Gradient Boosting Classifier | 0.758 | 0.776 |
| **Merged Descriptors** |  |  |
| Support vector classifier | 0.653 | 0.689 |
| Random Forest | 0.717 | 0.711 |
| Random Forest(BorutaPy) | 0.679 | 0.678 |
| Gradient Boosting Classifier | 0.741 | 0.702 |
| Ensemble(RF+SVC) | 0.735 | 0.667 |

In the end, we also looked closely to see if there are any similarities in the samples that are being misclassified across all the models. We found that the samples in the test data set that were being misclassified in all the models across the three types of descriptors could be grouped into the following two categories.

1. Samples labelled attractant or neutral, all belonging to the study carried out by Bargmann et al [10]: There are 8 such samples for Seq2seq descriptors, 3 for RD-kit descriptors and 7 for merged descriptors. This could possibly be a reflection of threshold values used to define category boundaries in that particular study. This can be mitigated by using (i) uniform behavioral assays and (ii) a three class classification subject to availability of more data. To further verify this, we performed a multi class classification on the augmented one hot encoding data (Supplementary Figure 1), and found that the distribution of misclassifications from the confusion matrices point towards the model performing poorly on classification between attractants and neutral which might be an artifact of threshold boundaries.
2. Ions: Only the RDkit descriptors set had ions in the dataset and three of them - all labelled repellants, were misclassified in all the models.

Despite a few misclassifications, this analysis shows that there is a non-trivial relationship between the arrangement of atoms that make up the chemical and the chemosensation and behavior exhibited by *C. elegans*.

### B. Training on an augmented data set leads to better accuracy

Having established the existence of the structure behavior relationship between the odorant and the organism, we next sought to increase the accuracy of the models. One way to do this is to have a larger data set for training. For this, we converted the SMILES format of the chemical into one-hot encodings and applied the augmentation methodology developed by Bjerrum [71]. Table II summarises the F1 scores for the models using one-hot encodings as features for both the original and the augmented data sets. SVC, RF and ensemble models give comparable scores in both the random states. However, scores are higher when training is carried out on the augmented data set instead of the original data set for all the models. The AUC-ROC curves for the ensemble models in both the random states, shown in (Fig.4), depict this increase. One can also notice that these curves are smoother for the augmented data set because of the increased number of samples that the model can be trained on.

**Figure 4:**
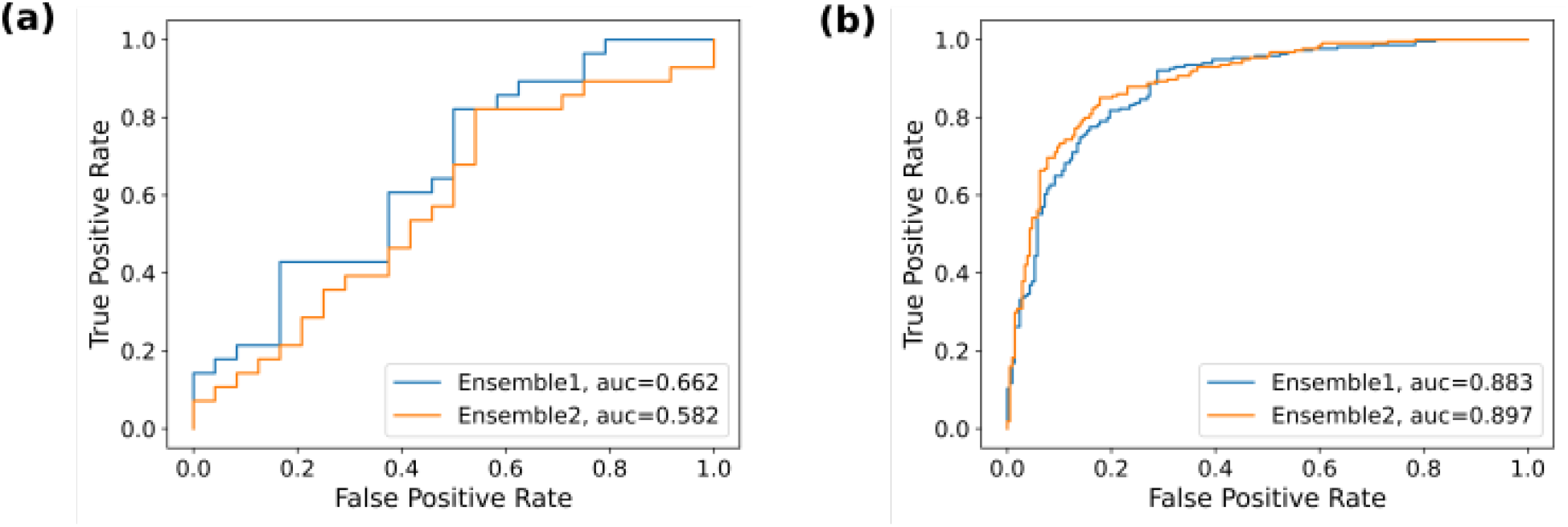
AUC-ROC curves for ensemble models for the first random state (blue) and the second random state (orange) for (a) the original and (b) the augmented data set.

**Table II:** F1 scores of various models

| Model | Random state 1 | Random state 2 |
| --- | --- | --- |
| **one-hot encodings** |  |  |
| Support vector classifier | 0.645 | 0.667 |
| Random Forest | 0.667 | 0.687 |
| Ensemble | 0.620 | 0.704 |
| **one hot encodings - augmented** |  |  |
| Support vector classifier | 0.796 | **0.835** |
| Random Forest | 0.816 | 0.761 |
| Ensemble | 0.815 | **0.839** |

Finally, deep neural networks, specifically Long Short-Term Memory (LSTM) networks, like the one used by Bjerrum, were trained on the vectorised data both by automated augmentation using data generators and the augmented data set produced earlier. In both the cases, deep neural networks failed to match the performance of RF and SVC models used on the original data set containing 173 entries. Therefore, we did not pursue this methodology further. The most probable reason for the failure of the LSTM model on this particular data set is the data hungry nature of neural networks. This causes the model to remain under trained in the absence of a data set much larger than what was achieved via augmentation.

## IV. DISCUSSION

We began our analysis by splitting the data into a test and training set using two different random states. Following this, we fit the data to simple RF and SVC models. We found that both the RF and SVC models in the two random states were overfitting to the training data, and in order to avoid that, we used principal component analysis onto our data. This reduced the overfitting and gave better scores with an average improvement of 3.9 % for RF and 0.8 % for SVC, respectively. To improve the models further, we tuned the hyperparameters using grid search and five fold cross validation implemented through the GridSearchCV method of the Sci-kit learn library. The hyperparameters thus obtained were used to build models that were more robust and less prone to overfitting while giving comparable or better F1 scores. We further used a voting classifier to take a vote of the two models obtained after the hyperparameter tuning. This adds up the probability scores from the given models and gives out a result with the highest sum of probabilities. As the data was split using two random states, we obtained two voting classifiers, and the average of the two F1 scores (0.8) was then used to compare with the standalone models. The four models - two RFs and two SVCs all have comparable scores to the average of the two ensemble models. However, the two voting classifiers give better results with a high F1 score (0.800) (Table I) and can be selected from all. We also trained other models like KNN and GBC on the same data and compared the scores to that of SVC, RF and the voting classifier. Both performed comparably to the previous models ( Avg F1 scores: KNN = 0.765, GBC = 0.800), with GBC outperforming all the other models.

We repeated the process for RDKit (Avg F1 score= 0.714) and Merged (Avg F1 score= 0.697) descriptors and found that Seq2seq descriptors (Avg F1 score= 0.77) performed better than the other two. Here too, the GBC gave comparable scores to the voting classifiers and outperformed all other models. The average F1 score seems restricted by limited data availability and increased from 0.665 to 0.81 when an augmented data set was used with ensemble models trained on one-hot encodings of the same.

Our analysis gives consistent prediction scores and points towards the presence of an underlying relationship between odorant structure and worm behaviour. However, it suffers from the drawback of not being able to differentiate between enantiomers, which, even though they have the same SMILES string, are known to smell different [74]. Additionally, the data used is only for monomolecular olfactants as compared to the complex mixture of odors an organism encounters in it’s natural environment. Further, because of a lack of consensus about a single standard experimental method in the scientific community, the data itself is taken from multiple sources that used different concentrations of odorants and different measurements to determine if the worm is attracted to an odorant or not. However, since we are limited by data from the literature, as mentioned in Section II, we have removed all conflicting or contradictory samples from the data set, thereby ensuring at least some degree of uniformity.

Another shortfall is the interpretability of descriptors. The Seq2seq descriptors don’t correspond to any physico-chemical properties and cannot tell us what property of the chemical structure is important for predicting worm behaviour. Moreover, the use of PCA as a dimensionality reduction method compromises the interpretability of features. To shed some light on the importance of features, we have used Boruta (listed as BorutaPy in Table I) for feature selection as an alternative to PCA. In the case of RDKit descriptors, where the features correspond to physico-chemical properties, we found that‘Chi1n’, ‘Kappa1’, ‘MinAbsPartialCharge’ were selected for the first random state and ‘Kappa1’, ‘MaxAbsPartialCharge’ for the second random state. The ‘MinAbsPartialCharge’ and the ‘MaxAbsPartialCharge’ return the minimum and the maximum absolute charge, respectively, for the molecule considering all the atoms. ‘Chi1n’ and ‘Kappa1’, on the other hand, are molecular graph theory based descriptors corresponding to first order molecular connectivity chi index and first order molecular shape attribute respectively [75]. The emergence of these descriptors reflects upon the significance of the chemical structure and the constituent atoms. However, a more detailed study will require a larger data set than currently available.

In order to improve upon and further validate the analysis, experimental methods involving a standardised chemotaxis assay can be employed for a set of chemicals, and numerical chemotactic index values can be calculated and fed into machine learning models in place of the current categorical binaries. It has been shown that the incorporation of odorant receptor data increases the accuracy of prediction in humans [76]. However, there is only a limited set of olfactory receptor data available for *C. elegans* due to the complexity arising from their co-expression on neurons and the presence of a large number(≈1200) of GPCR genes [30, 77]. As and when more data is available, that too can be incorporated into the model. Alternatively, simulated olfactory receptor data as generated by Milanetti et al. [30] can be used for the same. Further, with the neural connectome being complete, more advanced models incorporating neuronal activity and network connectivity can be constructed.

Finally, this study concludes that there exists a non-trivial relationship between the odorant and the olfactory behavior that can be harnessed to predict the action of worms in response to a given odorant. With ongoing research on odorant receptor characterisation and that on deciphering the underlying circuitry [78], *C. elegans* can serve as a model system for a detailed study of olfactory systems and help crack the olfactory code.

## DATA AVAILABILITY

The data that supports the findings of this study are available within the article.

## Notes

### Competing Interest Statement

The authors have declared no competing interest.

