## Supplemental text for "A machine learning based analysis to probe the relationship between odorant structure and olfactory behaviour in *C. elegans*"

***Multi class Classification:*** We generated one hot encodings of the multiclass data with the classes being "-1" for Repellent, "0" for Neutral and "1" for Attractant and generated an augmented data set as previously described. We further trained SVC and RF classifiers on it, the confusion matrices for which are given in Fig 1.

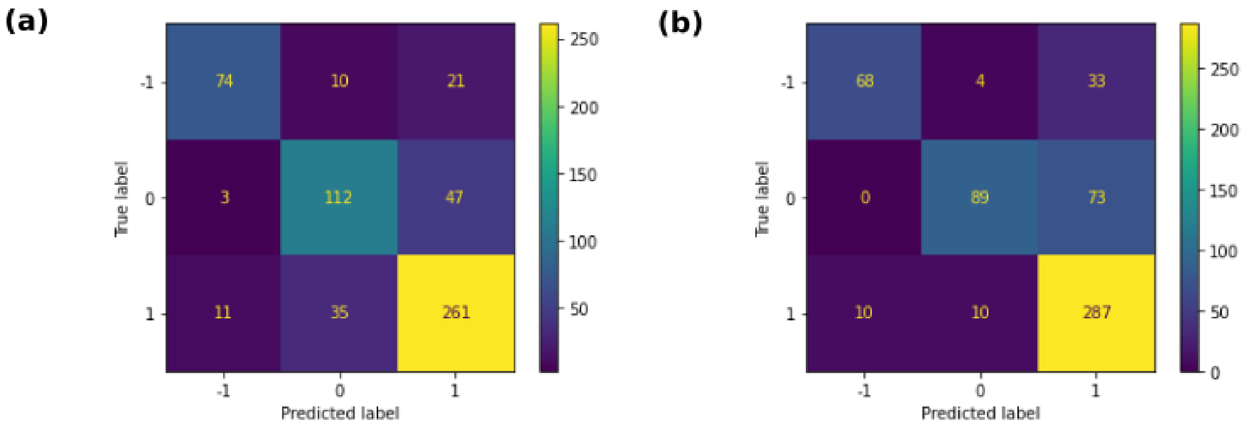


Figure 1: **Confusion Matrices for multi-class classification for** (a) Support Vector Classifier (b) Random Forest

*

**Table S1** The CSV file containing *C. elegans* response to odorants is provided.
**Table S2** F1 scores of the various models are given here.

**Seq2seq descriptors**

| **Model** | **Random state 1** | **Random state 2** |
|---|---|---|
| RF | 0.721 | 0.742 |
| RF PCA | 0.78 | 0.762 |
| RF BorutaPy | 0.733 | 0.7 |
| SVC | 0.754 | 0.706 |
| SVC PCA | 0.73 | 0.746 |
| SVC Tuned | 0.787 | 0.75 |
| Adaboost | 0.621 | 0.677 |
| Adaboost pca | 0.71 | 0.633 |
| KNN | 0.679 | 0.759 |
| KNN pca | 0.764 | 0.655 |
| KNN tuned | 0.764 | 0.767 |
| Ensemble (RF+SVC) | 0.8 | 0.8 |
| Ensemble with KNN | 0.75 | 0.787 |
| XGBoost | 0.69 | 0.698 |
| XGBoost pca | 0.691 | 0.69 |
| Gradient Boost | 0.759 | 0.73 |
| Gradient Boost pca | 0.807 | 0.797 |

**RDKit Descriptors**

| **Model** | **Random state 1** | **Random state 2** |
|---|---|---|
| RF | 0.716 | 0.689 |
| RF PCA | 0.725 | 0.746 |
| RF BorutaPy | 0.706 | 0.708 |
| SVC | 0.767 | 0.638 |
| SVC PCA | 0.75 | 0.714 |
| SVC Tuned | 0.75 | 0.633 |
| Adaboost1 | 0.656 | 0.727 |
| Adaboost pca | 0.708 | 0.696 |
| KNN 1 | 0.677 | 0.699 |
| KNN pca | 0.698 | 0.644 |

| | | |
|---|---|---|
| KNN tuned | 0.698 | 0.656 |
| Ensemble (RF+SVC) | 0.758 | 0.698 |
| Ensemble with KNN | 0.698 | 0.656 |
| XGBoost | 0.727 | 0.719 |
| XGBoost pca | 0.646 | 0.696 |
| Gradient Boost | 0.794 | 0.754 |
| Gradient Boost pca | 0.758 | 0.776 |

**Merged Descriptors**

| **Model** | **Random state 1** | **Random state 2** |
|---|---|---|
| RF | 0.72 | 0.618 |
| RF PCA | 0.717 | 0.712 |
| RF BorutaPy | 0.679 | 0.678 |
| SVC | 0.679 | 0.667 |
| SVC PCA | 0.679 | 0.688 |
| SVC tuned | 0.653 | 0.688 |
| Adaboost | 0.64 | 0.633 |
| Adaboost pca | 0.64 | 0.712 |
| KNN 1 | 0.654 | 0.633 |
| KNN pca | 0.654 | 0.656 |
| KNN tuned | 0.641 | 0.633 |
| Ensemble (RF+SVC) | 0.735 | 0.667 |
| Ensemble with KNN | 0.654 | 0.688 |
| XGBoost | 0.612 | 0.645 |
| XGBoost pca | 0.72 | 0.69 |
| Gradient Boost | 0.764 | 0.698 |
| Gradient Boost pca | 0.741 | 0.702 |

**One hot encodings**

| **Model** | **Random state 1** | **Random state 2** |
|---|---|---|
| RF | 0.604 | 0.576 |
| RF PCA | 0.667 | 0.687 |
| SVC | 0.667 | 0.571 |
| SVC PCA | 0.656 | 0.693 |

| | | |
|---|---|---|
| SVC Tuned | 0.645 | 0.667 |
| Adaboost | 0.468 | 0.586 |
| Adaboostp pca | 0.641 | 0.635 |
| KNN | 0.608 | 0.586 |
| KNN pca | 0.571 | 0.615 |
| KNN tuned | 0.645 | 0.536 |
| Ensemble (RF+SVC) | 0.62 | 0.704 |
| XGboost | 0.536 | 0.552 |
| XBboost pca | 0.63 | 0.613 |
| Gradient Boost | 0.566 | 0.657 |
| Gradient Boost pca | 0.646 | 0.714 |

| | **One hot encodings-Augmented** | |
|---|---|---|
| **Model** | **Random state 1** | **Random state 2** |
| RF | 0.811 | 0.822 |
| RF PCA | 0.816 | 0.761 |
| SVC | 0.811 | 0.799 |
| SVC PCA | 0.798 | 0.812 |
| SVC Tuned | 0.795 | 0.835 |
| Adaboost | 0.686 | 0.732 |
| Adaboost pca | 0.718 | 0.648 |
| KNN | 0.757 | 0.752 |
| KNN pca | 0.72 | 0.773 |
| KNN tuned | 0.737 | 0.757 |
| Ensemble (RF+SVC) | 0.815 | 0.839 |
| XGboost | 0.78 | 0.813 |
| XBboost pca | 0.803 | 0.757 |
| Gradient Boost | 0.638 | 0.663 |
| Gradient Boost pca | 0.684 | 0.71 |

Table S2: F1 Scores
